# The trans-regulatory landscape of gene networks in plants

**DOI:** 10.1101/2022.10.23.513368

**Authors:** Niklas F. C. Hummel, Andy Zhou, Baohua Li, Kasey Markel, Izaiah J. Ornelas, Patrick M. Shih

## Abstract

The effector domains of transcription factors play a key role in controlling gene expression; however, their functional nature is poorly understood, hampering our ability to explore this fundamental dimension of gene regulatory networks. To map the trans-regulatory landscape in a complex eukaryote, we systematically characterized the putative effector domains of over 400 *Arabidopsis thaliana* transcription factors for their capacity to modulate transcription. We demonstrate that transcriptional effector activity can be integrated into gene regulatory networks capable of elucidating the functional dynamics underlying gene expression patterns. We further show how newly characterized domains can enhance genome engineering efforts and reveal how plant transcriptional activators share regulatory features conserved across distantly related eukaryotes. Our results provide a framework to systematically characterize the regulatory role of transcription factors at a genome-scale in order to understand the transcriptional wiring of biological systems.

## Main Text

Biological systems are reliant on transcriptional networks, which are largely regulated by transcription factors (TFs). At their core, TFs are defined by two broad functions: 1) specific binding of target cis-regulatory DNA sequences through DNA-binding domains (DBDs) and 2) regulating transcription (i.e., gene activation or repression) through their transcriptional effector domains (TEDs). TEDs can serve as biochemical beacons recruiting or inhibiting transcriptional machinery; however, the mechanisms underlying these processes are not well understood and have primarily been studied in other eukaryotes distantly related to plants (i.e., yeast, human, etc.)^1^. Recent technical advances and large consortium efforts have dramatically expanded our understanding of TF binding sites across full genomes^2,3^. However, the nature of these interactions has remained elusive, as the functional characterization of TEDs has not been as readily scalable. As a result, our knowledge of TEDs that compose the trans-regulatory landscape has not kept pace with the characterization of its cis-regulatory counterpart^2,4,5^. Hence, identification and characterization of these domains in plants is an important first step towards elucidating the design principles that govern gene regulation in order to ultimately enable more refined approaches to engineer and fine-tune transcription in plants.

Unraveling the functional dynamics of gene regulatory networks (GRNs) is a key challenge of systems biology with the promise to understand the regulatory architecture of biological systems. To observe how genome-scale regulation of transcription occurs, GRNs hinge on the either activating or repressing interactions between individual TFs and their target genes. Hence, a central goal of the field of systems biology is to map genome-scale GRNs to understand the concerted regulation of biological phenomenon and traits^6,7^. However, due to the lack of knowledge of the regulatory role of TFs, GRNs are largely limited to TF binding site and RNA-seq based co-expression information which can only indirectly infer whether TF-gene interactions are activating or repressing^6,8^. Therefore, we reasoned that directly measuring the regulatory function of TEDs and integrating this information into GRNs could provide a missing, yet integral, dimension to studying the underlying regulatory wiring of biological systems.

Modulating the expression of plant genes has been a key area of focus for precision crop engineering, as many agronomically important traits are the result of altered gene expression ^9,10^. Hence, the intrinsic trans-regulatory elements embedded in plant TF proteins offer a unique resource to mine for novel TEDs that may advance plant engineering efforts and understanding their native regulatory role in GRNs could provide novel targets for engineering. To expand our understanding of plant transcriptional regulation, we systematically measured the activation or repression activity of putative TEDs from over 400 *A. thaliana* TFs, providing unique insights into the underlying biochemical properties of plant TEDs. We further show how the integration of trans-regulatory information into GRNs can validate and describe the functional role of TFs in gene networks. Our findings demonstrate how genome-wide functional characterization of TEDs can enhance our understanding of the transcriptional regulation of biological systems, both on a biochemical and systems level.

## Results

### Genome-wide characterization of plant transcriptional effector domains

The *in vitro* DNA binding activity of 529 *A. thaliana* TFs has been previously reported^2^; however, mapping TF-DNA interactions alone cannot provide information on the regulatory nature of these interactions, limiting our ability to understand key facets of plant gene networks. Previous attempts in large-scale characterization of TEDs in human and yeast models have focused on short length TEDs (≤80 amino acid)^11,12^. While such studies yield short peptides with transcriptional effector activity, they may not fully capture the regulatory activity of the TF. As an example, some activators rely on multiple long subdomains for activity^13^. Thus, in order to provide a more comprehensive understanding of TF function, we instead focused on experimentally characterizing size unrestrained TEDs of *A. thaliana* TFs whose DNA binding motifs and downstream targets have previously been mapped^2,14^. For characterization of TEDs, we utilized a transient, synthetic transcriptional system in *Nicotiana benthamiana* that we previously established^14^. First, we generated putative TEDs by identifying and excluding conserved DBDs and selectively extracted the longest non-DBD TF protein sequences which ranged from 27 to 779 amino acids (Fig. S1). We then fused these candidate TEDs to the yeast Gal4 DBD, generating a library of synthetic TFs (SI Table 1). The Gal4 DBD localizes the TED candidate to a plant synthetic reporter composed of a minimal promoter with 5 concatenated Gal4 binding sites driving GFP^14^. By measuring GFP fluorescence in the presence of a synthetic TF and normalizing the signal for basal expression of the reporter by using a constitutively expressed dsRed, we can individually characterize the functional role of TEDs independent of their regular protein context (Fig. 1A).

**Figure 1.**
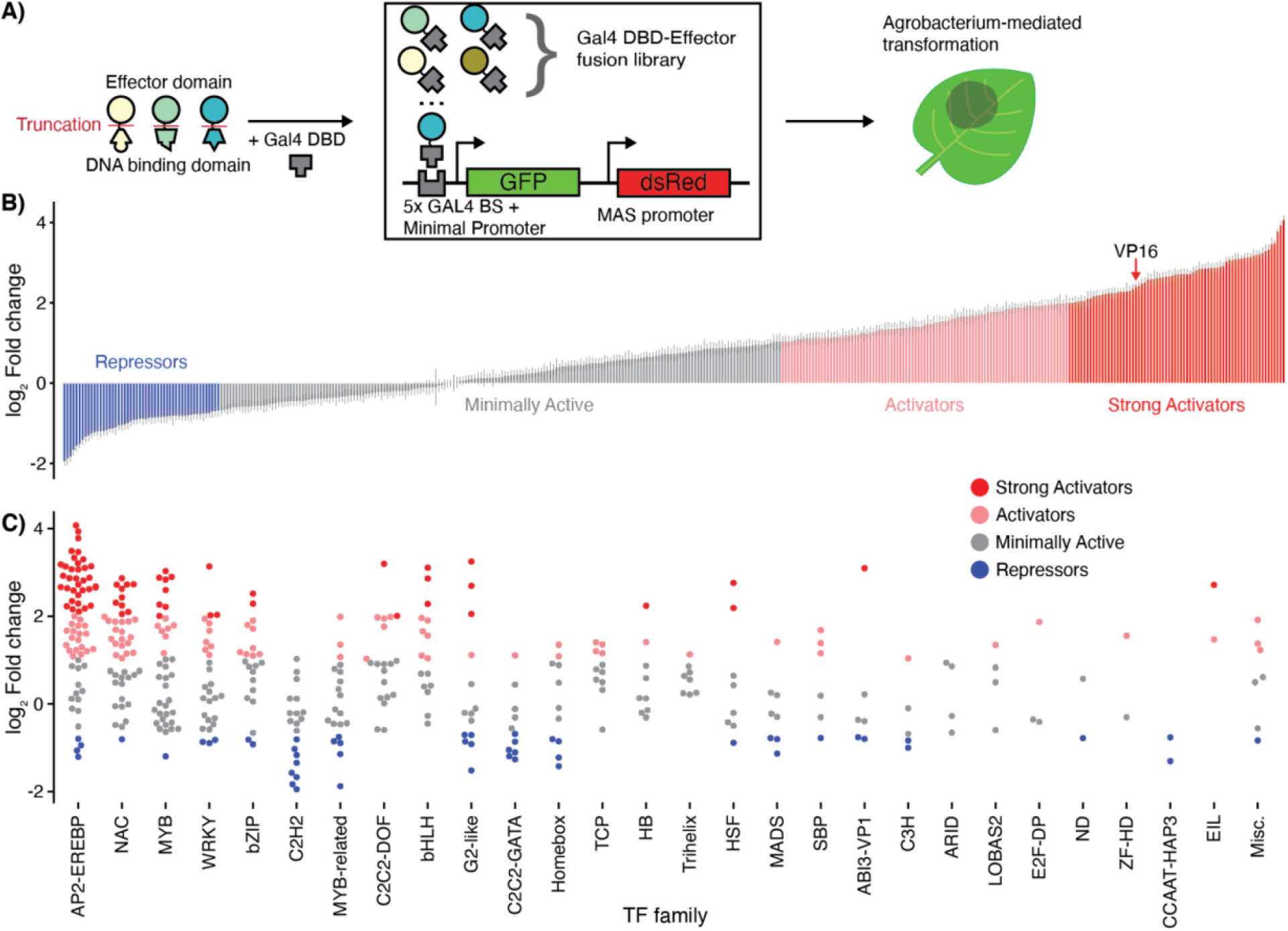
Genome-scale characterization of hundreds of plant transcriptional effector domains. (**A**) Truncated putative TEDs are fused to the yeast Gal4-DBD to generate a library of synthetic TFs. Gal4-TED fusions bind to a synthetic promoter and their effect on transcription is measured via a fluorescent reporter GFP and normalized using a constitutively expressed dsRed. (**B**) Normalized GFP expression of 403 synthetic TFs in relation to background reporter expression in *N. benthamiana* leaves 3 days post infiltration (n=16 biological replicates). Arrow indicates the position of Gal4-VP16 as a strong activator control. (**C**) Normalized GFP expression from (B) grouped by TF family. Individual data points represent single TEDs characterized in this study. TF families with single TEDs in the screen were grouped into Misc. and include REM, LIM, RAV, zf-GRF, RWP-RK, BSD and NLP.

Using this approach, we individually characterized 403 synthetic TFs using transient expression in *Nicotiana benthamiana* (Fig. 1B, SI Table 2). We identified 166 activator and 53 repressor domains, defining activation and repression as an increase of GFP expression by at least 100% and decrease by at least 40% in comparison to basal expression of the reporter based on statistical thresholds (see Methods). We found 49 activators displaying stronger trans-activation activity than the widely used viral activator VP16, with the strongest activator derived from the cold response TF CBF4 increasing GFP expression 16-fold over basal reporter expression (Fig. 1B). Our findings demonstrate the potential of transient gene expression in *N. benthamiana* to systematically study TEDs and enable the development of enhanced genetic engineering tools, providing alternatives to broadly used TEDs like VP16.

In order to validate the findings of our assay, we compared our observed TED activity with the activity of each parent TF as a repressor, an activator, or both according to previous studies. We found a large overlap between the activity of TEDs in this study and TFs individually studied *in vivo* (SI Table 3). Of our 166 annotated TEDs with activator activity, 90 have been previously reported to directly activate expression, and only 7 act as repressors, with 1 as both an activator and repressor. Of our 52 TEDs with repressor activity, 21 have been previously shown to act as repressors, 2 as activators, and 1 as both an activator and repressor. Notably, we characterized 68 novel activators and 28 novel repressors, highlighting how our approach enables the discovery and characterization of new TEDs. The broad overlap of referenced TF activity with the TEDs described here indicates the consistency and transferability of TED activity between our synthetic heterologous system and *in vivo* observations in the native plant. To further validate our repressor findings, we studied the occurrence of the well-known repressive EAR motif in our TED populations. As expected, we found an overrepresentation of the repressive EAR motif in the repressor population (33% of TEDs with motif) when compared both to the activator (4%) and minimally active populations (13%, Fig. S2), supporting the agreement between our assay and the *in vivo* function of TEDs in their natural context. Taken together, we found a large qualitative agreement between our TED characterization and published *in vivo* activity which enabled us to combine DNA binding and TED activity for further analysis.

Our dataset spans TEDs from 34 TF families and allows us to study functional trends across TF families (Fig. 1C). For example, the AP2-EREBP TF family comprises 147 TFs in *A. thaliana* and of the 70 AP2-EREBP TFs studied here only 4 TEDs act as repressors and 54 TEDs as activators (Fig. S3). This indicates a bias towards activators inside the AP2-EREBP family. Conversely, in the C2H2 TF family we found 8 out of 20 TEDs studied here to significantly repress gene expression with none as characterized activators. These observations overlap with human C2H2-TFs which mostly contain repressive TEDs^15^. Thus, regulatory roles across long evolutionary distances might be conserved in the C2H2 family. TF binding sites within the same family are often redundant and general trends of specific TF families as either activators or repressors may increase the robustness of the regulatory network through a form of functional redundancy.

### Trends in TF downstream targets based on TED activity

Combining TF binding site and TED activity information allowed us to study trends in which TF-gene interactions are either repressive or activating. By incorporating this regulatory logic, it becomes possible to describe recurring network motifs which are broadly found across biological systems^16^. One such network motif in prokaryotes is negative autoregulation (NAR), where a repressor downregulates its own expression^17^. NAR enables the acceleration of response times and reduces cell-to-cell variation in protein concentration thus enabling robust regulation of their targets^16,18^. To investigate usage of NAR in plant TFs, we combined TED activity with published DNA binding data^2^. When comparing TED populations based on their regulatory activity, we found that repressors were more likely to auto-regulate than strong activators (defined as TEDs increasing gene expression more than 400%, Fisher’s exact test P = 0.04, Fig. S4A). We also searched for a bias between positive and negative feedback loops, i.e., two TFs regulating each other, but did not observe any bias between the activator and repressor populations (Fig. S4B). Our analysis of NAR in plants supports convergence of these motifs across both prokaryotes and eukaryotes supporting previously suggested emergent properties of transcriptional regulation.

We further identified trends in the functional roles of genes targeted by TFs with strongly activating TEDs. We analyzed the gene ontology (GO) terms of genes targeted by activators stronger than VP16. Interestingly, we found that the GO terms of these genes were enriched for terms linked to response to hormones, stresses, and external stimuli. GO terms linked to primary or secondary metabolism were depleted, except for terms linked to cell wall biogenesis (Fig. S5). This is consistent with a requirement for strong gene activation to enact the rapid changes to transcriptional programming needed for a concerted response to stimuli rather than the direct activation of metabolic pathway genes during housekeeping functions.

### An integrated cis- and trans-regulatory gene network elucidates the functional dynamics of TF-gene interactions

GRNs integrating cis-regulatory information and RNA-seq attempt to elucidate the underlying regulatory logic of TFs and their targets^8,19^; however, these GRNs are limited to indirect inferences of TF activity, rather than directly assaying how target TFs modulate gene expression. Thus, integrating trans-regulatory TED data as a layer on top of established GRNs could enhance their explanatory power and help study how TF-DNA interactions translate into regulatory output. As an ideal case study for this approach, we chose to investigate the well-characterized transcriptional response to nitrogen in *A. thaliana*^6^.

Recently, Varala *et al*. generated a high confidence nitrogen-responsive GRN encompassing 37 TFs and 171 direct genomic targets by combining published TF DNA binding with temporal RNA-seq^8^. Using our TED data, we annotated the links between TFs and their targets in this network as activating or repressing, thereby generating the first GRN integrating TED activity with DNA binding and temporal RNA-seq data (Fig. 2A, SI Table 4).

**Figure 2.**
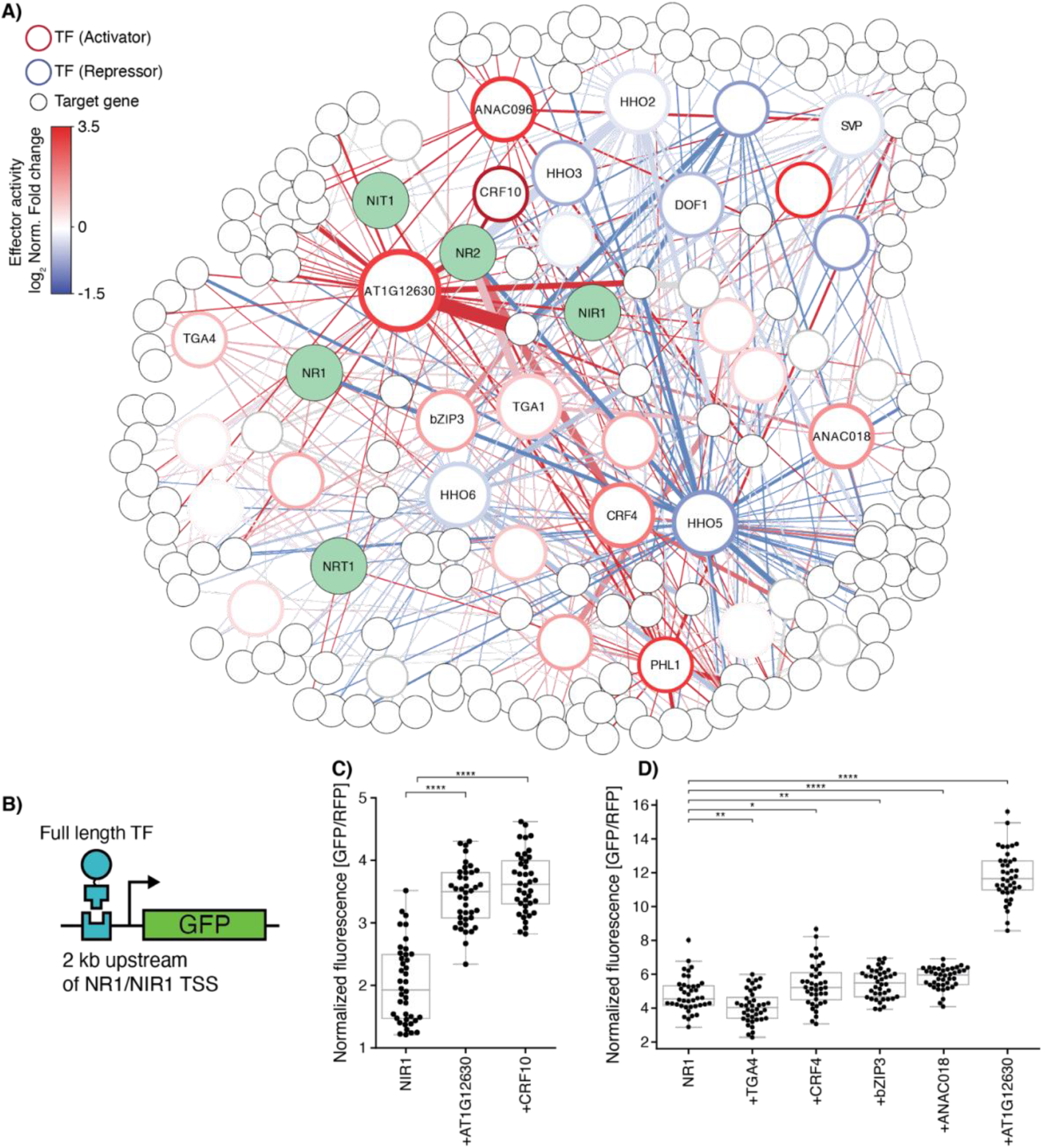
An integrated cis- and trans-regulatory GRN on nitrogen response in Arabidopsis. (**A**) GRN describing both TFs and respective target genes showing significant temporal changes in RNA abundance in response to nitrate in *A. thaliana*. Hollow nodes with colored edges depict TFs with respective trans-regulatory activity from this study. Target genes are small white nodes with black borders. Edges are based on experimentally verified DNA binding of TFs to target genes and annotated with TED activity data (color) and the predicted influence of a TF to its target (edge width)^8^. Green nodes indicate core nitrogen metabolism genes. (**B**) Coexpression of native full lengths TFs enables modulation of GFP expression driven by native NR1/NIR1 promoter regions. TSS: Transcriptional start site. Normalized GFP fluorescence for coexpressed full length activators with promoters derived from (**C**) NIR1 and (**D**) NR1. Asterisks indicate Mann-Whitney U test * P ≤ 5 × 10^−2^, ** P ≤ 5 × 10^−3^, **** P ≤ 5 × 10^−5^.

Annotating TF-gene interactions with regulatory activity relies on the assumption that the transcriptional effect of our Gal4-TED fusions on the reporter construct resembles the endogenous effect of its parent TF on its targets. We therefore sought to verify this hypothesis by testing if the activity of full length TFs is consistent with our measured TED activities in a synthetic transcriptional system. We reconstituted native TF-DNA interactions of core nitrogen metabolism genes by building GFP reporter constructs driven by the *Arabidopsis* promoters of nitrate reductase 1 (NR1) and nitrite reductase 1 (NIR1) (Fig. 2B). We then co-expressed full length TFs that are predicted to regulate these genes throughout the entire time course according to the GRN. We found that the TFs CRF10 and AT1G12630 – both annotated as activators in our assay and predicted to interact with NIR1 – significantly increased the activity of the NIR1 promoter (Fig. 2C). Five TFs whose TEDs are activators in our assay – CRF4, bZIP3, TGA4, ANAC018 and AT1G12630 – are predicted to interact with the NR1 promoter in the GRN. All five TFs altered expression with AT1G12630 strongly inducing, CRF4, bZIP3 and ANAC018 weakly inducing, and TGA4 repressing the NR1 promoter. CRF4, bZIP3 and TGA4 have been previously shown to induce gene expression during nitrogen response^20^. Hence, we were able to validate the regulatory activity of 6 out of 7 TF interactions with core nitrogen promoters. The repression of transcription by TGA4 may be a result of lacking co-activators, or secondary mechanisms that are found in the native system. Nevertheless, the fact that our heterologous system can elucidate activity by observing single TF-promoter interactions confirms that gene regulatory networks can be enriched by annotating TED activity.

To further validate that TFs in GRNs can be annotated as activators or repressors based on TED activity, we sought to exploit the temporal dynamics of target genes in the nitrate response network by Varala *et al*. ^*8*^. For example, genes targeted by TFs whose TED is an activator in our assay should tend to increase in mRNA abundance over time. By leveraging the temporal nature of the Varala *et al*. dataset, we validated our measured TED activities and observed the causal relation between TF and the transcriptional output of their downstream targets. Nitrate responsive genes show altered gene expression early after nitrate induction^21^. Therefore, we focused on the early nitrogen response between 0-30 min. At 15 min post nitrate induction, we observed a set of six annotated activators which target primary nitrate response genes (NR1, nitrate reductase 2 (NR2), and NIT1) (Fig. S6A). The presence of activators should lead to induction of target genes. NR1/2 and NIT1 indeed show increased levels of expression at later time points (Fig. S6B) with increased expression after the interaction with our characterized activators including the validated bZIP3 and AT1G12630. We further observed the RNA abundance of all genes targeted by this activator group (Fig. S6C). As a measure for dynamic changes in gene expression we calculated the rate of expression change in between every time point for every gene of this group (Fig. S6D). As expected, we found that between 20 and 30 min the majority of genes targeted by the activators present in the 15 min sub network showed their largest rate of expression induction (Fig. S6E), and none showed their strongest reduction of expression (Fig. S6F). This indicates that nitrate responsive genes targeted by this activator group show the predicted response based on trans-regulatory activity *in vivo*.

Together our results demonstrate how integrating TED activity from heterologous experiments into GRNs can recapitulate the regulatory relationships of TFs with their downstream targets. Hence, systematic TED characterization provides an important means to fill in major gaps in our knowledge of GRNs that top-down observations have been unable to resolve.

### Novel plant activators expand modularity of synthetic biology and genetic engineering tools

Having shown that TED activity can enrich GRNs, we applied our novel TEDs in a synthetic biology context to control gene expression and expand the dynamic range of native gene transcriptional profiles. Previously developed plant synthetic biology tools have heavily relied on a small subset of characterized effectors. For example, the herpes simplex virus-based VP16 domain has been utilized in many eukaryotic systems, including plants, as the state-of-the-art activator since its discovery over 30 years ago^22,23^. Thus, it is of note that many of our characterized TEDs demonstrate stronger activation activity than the VP16 domain which is commonly used in genome engineering approaches (e.g., dCas9-based CRISPR activation, synthetic transcription factors, etc.)^24,25,26^. Our findings demonstrate how TED screens enable a powerful approach to mine for host-specific (e.g., plant-specific) activator domains that can be superior to the state-of-the-art domains currently utilized.

To explore the utility of newly discovered activation domains for metabolic and genome engineering, we tested how 20 activator domains stronger than VP16 would perform when fused to other TFs as a means to enhance their transcriptional output. We fused our chosen activator domains to the anthocyanin master regulator PAP1 as it activates the expression of multiple anthocyanin pathway genes resulting in a quantitative readout via elevated levels of anthocyanins in plant tissue (Fig. 3A)^27,28^. We expressed PAP1-activator fusions in *N. benthamiana* for 3 days and quantified the anthocyanin content by absorbance measurements of extracts from leaf tissue. Multiple activators showed increased concentration of anthocyanins in comparison to PAP1 and a PAP1-VP16 fusion (Fig. 3B, C). Of 20 activator candidates, 16 displayed higher absorbance values than PAP1 and 11 performed better than the PAP1-VP16 fusion (SI Table 5). Interestingly, the strongest activator TED from the screen, derived from CBF4, achieved the strongest increase in anthocyanin concentration. Taken together our panel of top activation domains can be easily integrated into synthetic biology applications like the optimization of the transcriptional output of TF master regulators.

**Figure 3.**
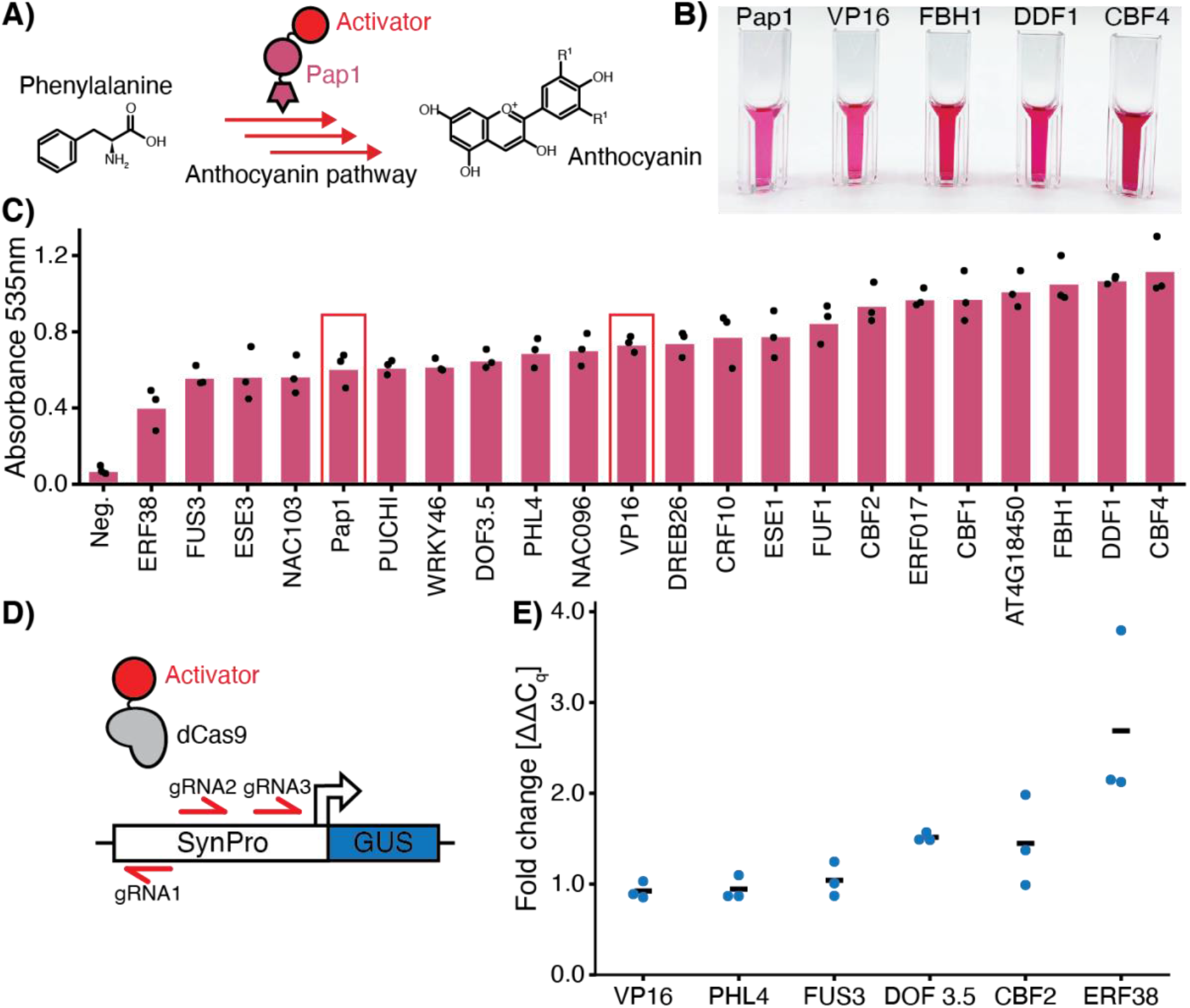
Strong plant activators outperform VP16 when integrated into synthetic biology applications. (**A**) Fusion of strong activators to the anthocyanin master regulator PAP1 promotes production of anthocyanins. (**B**) Visual representation of anthocyanin extracts quantified in (C). (**C**) Quantification of anthocyanins extracted from *N. benthamiana* leaf tissue expressing PAP1-fusion constructs. Red bars indicate Pap1 as a basal expression and Pap1-VP16 as a positive control. Neg.: Leafs infiltrated with no vector control. (**D**) Activator fusion to dCas9 to modulate target gene expression. Three gRNAs localize the dCas9-activator fusion to a synthetic promoter driving GUS. (**E**) Fold change in transcript abundance of dCas9-activator fusions relative to the reporter construct alone quantified by qRT-PCR.

The modularity afforded by deactivated RNA-guided nuclease variants (e.g., dCas9) allows for the targeted alteration of gene expression when selectively defined by engineered guide RNAs^25,29^. Thus, the versatility of the DNA binding capability of dCas9-activator constructs has been leveraged to enable genome wide CRISPR activation screens, but again have mostly relied on VP16-based viral activators. Hence, we sought to benchmark our top activator candidates against VP16 in an established expression system^30^. We fused five strong activators from our screen to dCas9 and compared these novel dCas9-activator fusions to dCas9-VP16 by targeting them to a synthetic promoter (Fig. 3D). We quantified transcript abundance by qRT-PCR with RNA extracted from *N. benthamiana* leaf tissue 3 days post *Agrobacterium*-mediated transformation. We observed that dCas9-VP16 displayed extremely low activity in comparison to two activator domains from ERF38 and DOF3.5 with ERF38 displaying significant increase of expression (Dunn’s test P < 0.005, Fig. 3E, SI Table 6); thus, our newly characterized domains have the potential to enhance CRISPR activation in plants. The field of genome engineering has embraced the use of VP16-based activators, and has largely coped with its low activation activity by recruiting large numbers of VP16 via various strategies^31,32^. As an alternative, our TED characterization demonstrates how identification of entirely novel, host-specific TEDs can result in an increased dynamic range of gene expression, and decrease reliance on TEDs that are not optimized to work in plants, such as VP16. Ultimately, our genome-wide screen enabled us to identify strong activator domains that can be used to tunably enhance transcription in a genome-specific manner, thereby providing a foundation for rapid generation of functional genomics toolsets.

### Conserved activity of plant transcriptional activators across eukaryotes

Just as the function of VP16 can cross eukaryotic super families^30,33^, plant transcriptional activation may utilize molecular machinery and mechanisms broadly conserved between distantly related species. In order to investigate the conservation of the regulatory activity of plant activators into other eukaryotes, we tested the ability of our twenty activators stronger than VP16 to promote constitutive gene expression in the model yeast system, *Saccharomyces cerevisiae*. We designed an expression cassette utilizing the well-characterized yeast inducible GAL1 promoter, which is induced in presence of galactose, repressed by glucose and contains Gal4 binding sites^34^, driving the fluorescent reporter GFP. We then observed the ability of Gal4-DBD-TED fusions to induce gene expression in the repressed state of the promoter using flow cytometry (Fig. 4A). TED activity was quantified by measuring the fractions of cells whose GFP expression was equal or higher than that of GAL1-GFP induced by galactose, while excluding observations similar to GAL1-GFP in glucose. When the Gal4-DBD-TED fusions were expressed constitutively, GFP expression was observed in <1% to 80% of the cell populations (Fig. 4A, SI Table 7). Notably, the TEDs derived from NAC103 and PHL4 were able to outperform VP16, marking them for further optimization in fungi (Fig. 4B). Importantly, the Gal4-DBD-activator fusions were tested in the presence of glucose, the repressed state of the GAL1 promoter. Still, multiple activators were able to enhance GFP expression, highlighting their potential for developing novel activation tools. Surprisingly, although some TF families like the AP2-EREBP TF family are plant-specific^35^, activators from this family function in yeast, suggesting that while evolved uniquely in plants, disparate TF families may have converged on similar mechanisms of activation.

**Figure 4.**
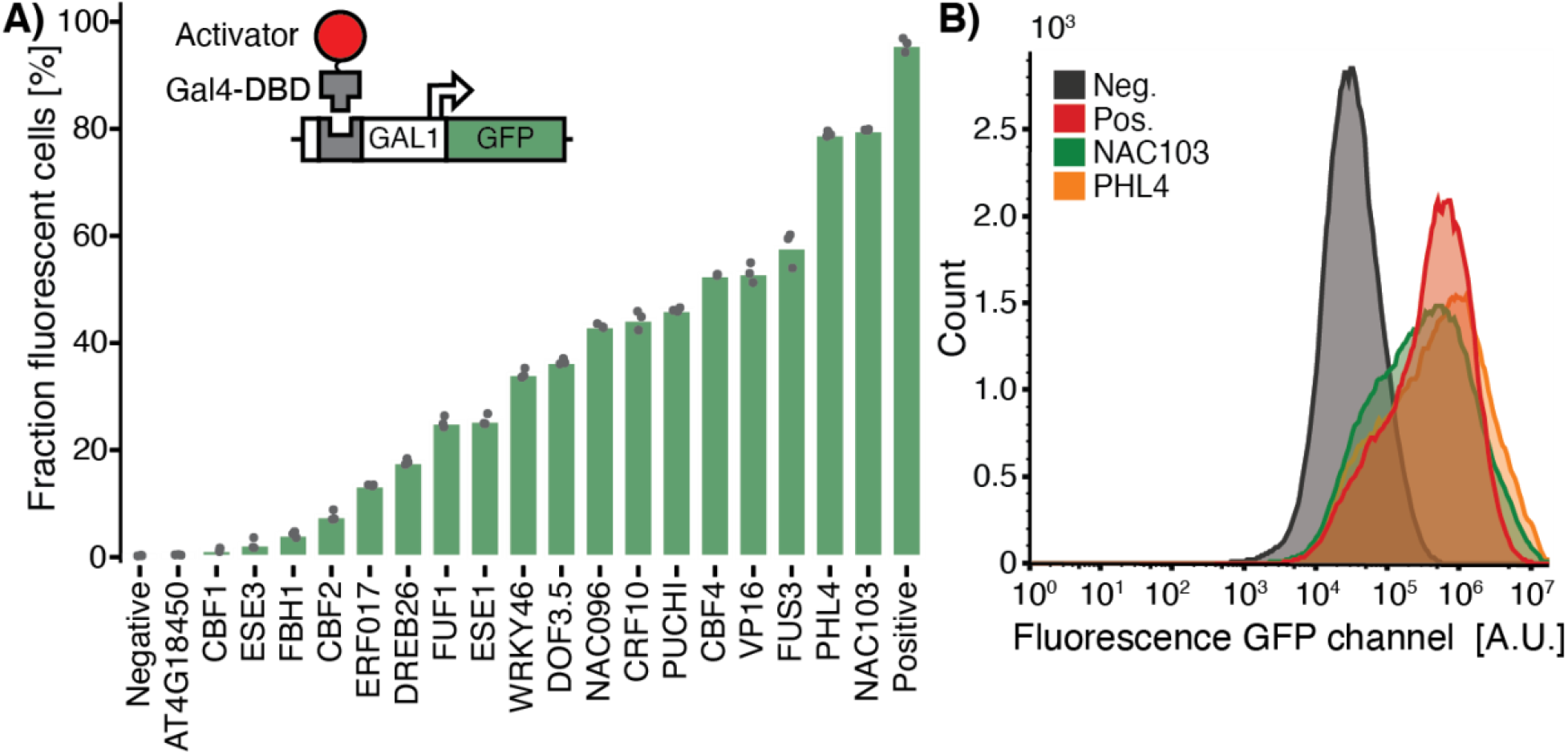
Plant activator activity is conserved in yeast. (**A**) Plant activators can induce a native yeast promoter when fused to the GAL4-DBD. Fractions of cells showing fluorescence in the repressed state of the GAL1 promoter grown in glucose. Each datapoint represents above threshold frequency for 100,000 recorded events. (**B**) GFP Fluorescence intensity distributions of activator and control populations for 100,000 recorded events.

The observation that plant activators can function in fungi suggested that there are conserved mechanisms underlying transcriptional activation between plants and yeast. We therefore analyzed the protein sequences of our TEDs using ADpred, a machine learning model trained on a large set of putative activation domains in 30 amino acid long protein sequences in *S. cerevisiae*^*36*^. We calculated the ADpred score for 30 amino segments of all TEDs in this study as described and assigned a binary value to every TED depending on whether it contained an amino acid section with an ADpred score ≥ 0.9. We found that activators are more likely to contain consecutive amino acid residues predicted to be activation domains than the repressor and minimally active populations (Fig. 5A, two-sided Fisher’s exact test, P = 0.00012). We found motifs in 19 out of our 20 chosen activators stronger than VP16 that scored above our ADpred threshold (Fig. S7). We extracted ADpred predicted subsections of three TED regions with strong activator activity (Fig. 5B), and benchmarked them against their full length TEDs and VP16 in *N. benthamiana* using the same assay as described in Fig. 1A. The ADpred predicted motifs of ESE3 and WRKY46 induced the expression of GFP similar to their full length TEDs and outperformed VP16 (Fig. 5C), showcasing the potential to mine plant TFs using the ADPred model. The two motifs of PHL4 were not able to induce GFP in the same manner as their parent TED suggesting that either the two motifs need to function as a bipartite motif or the parent TED uses a mechanism that the model cannot predict.

**Figure 5.**
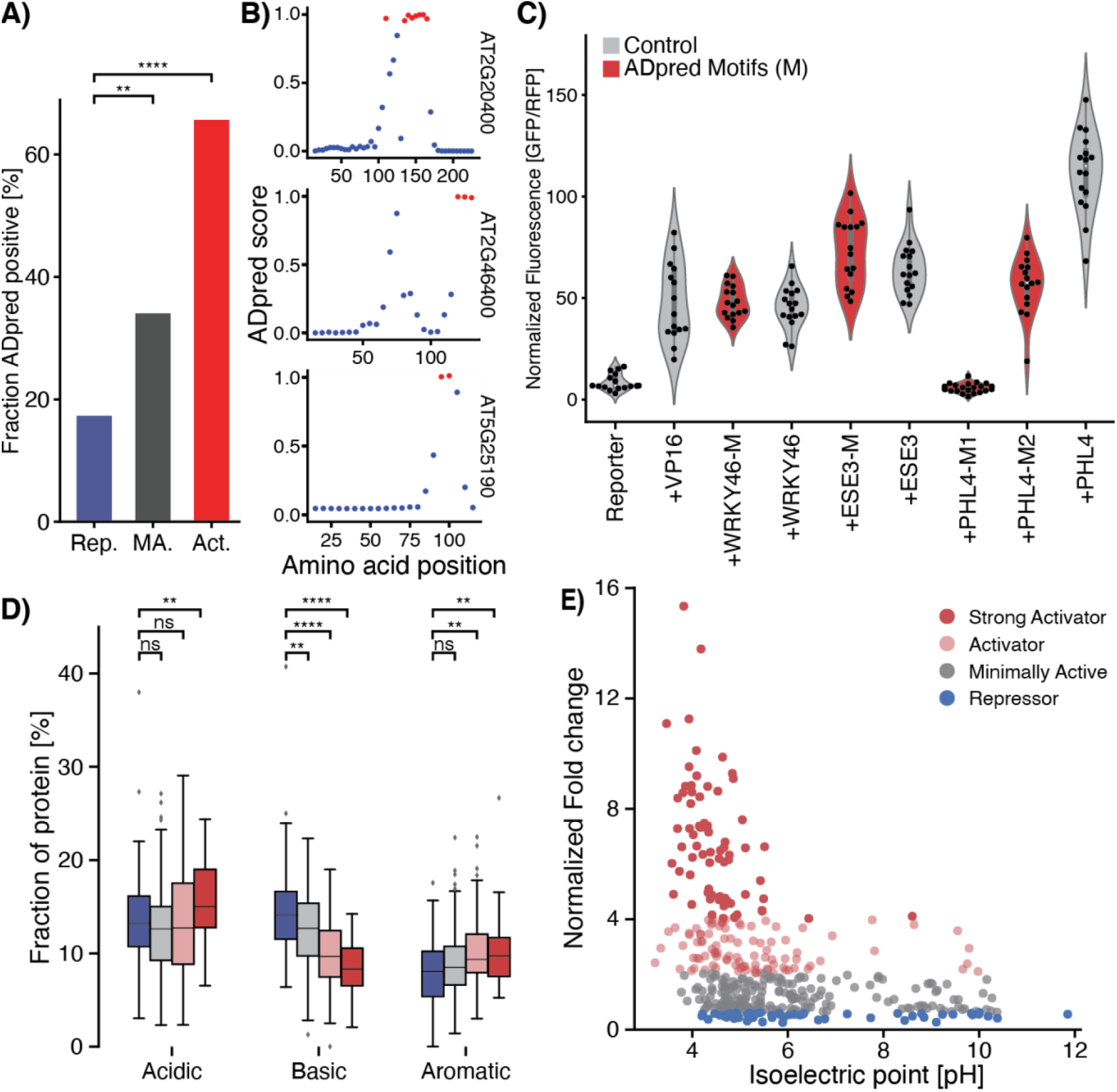
Translation of a yeast machine learning algorithm on plant TEDs supports trends in conserved biochemical features of activators. (**A**) Plant activators are enriched in activation domains predicted by the yeast machine learning model ADpred. Sum of TEDs with ADpred score ≥ 0.9 of each TED population. Asterisks indicate Fisher’s exact test ** P ≤ 5 × 10^−3^, **** P ≤ 5 × 10^−5^. Rep.: Repressor, MA. : Minimally Active, Act. : Activator. (**B**) ADpred analysis of three strong plant activators. ADpred scores were calculated for every 30 amino acid stretch slid along the protein sequence with window size = 5. Red dots indicate 30 amino acid long segments with ADpred score ≥ 0.9, blue dots < 0.9. (**C**) ADpred predicted activator motifs can perform similar to full length TEDs. Distribution of normalized GFP fluorescence reads for 16 biological replicates in *N. benthamiana*. Motifs found using ADpred indicated in red. M: Motif. (**D**) Amino acid frequencies of all individual candidate TEDs grouped into their respective population (asterisks indicate Mann-Whitney *U* significance test * P ≤ 5 × 10^−2^, ** P ≤ 5 × 10^−3^, *** P ≤ 5 × 10^−4^, **** P ≤ 5 × 10^−5^, ns non-significant). (**E**) Isoelectric point of TEDs mapped to performance in TED screen.

Because we demonstrate that machine learning models trained on yeast activation domains could reliably predict plant activation domains, we investigated if we could observe similarities in biochemical features between plant and yeast activators. Specifically, in yeast there have been well documented biases for acidic and large hydrophobic amino acid residues found in activation domains. We indeed found strong biases in the amino acid composition between the repressor and activator populations (Fig. 5D, Fig. S8A). Acidic, hydrophobic, and aromatic residues were significantly overrepresented among our characterized activators, whereas basic residues (e.g arginine, lysine and histidine) were significantly depleted and only enriched in repressors. These biases match the sequence profile of acidic activation domains found in mammalian and yeast systems^1,36–38^. The isoelectric point of TED populations also differed significantly, enforcing the importance of the amino acid composition. Activators show low isoelectric points, whereas repressors exhibit a wide range of isoelectric points. These data suggest that overall charge is a more important feature of activators than of repressors in plants (Fig. 5E). Structural disorder has also been suggested to be linked to TED activity, but, while TEDs were predicted to be on average >75% disordered, we did not observe a bias between TED populations (Fig. S8B). We further couldn’t find a bias between the protein length when comparing TED populations (Fig. S8C). Taken together our results indicate that plant activators share sequence features with their counterparts from distant eukaryotes suggesting the utilization of a general eukaryotic mechanism for transcriptional activation.

## Discussion

A holistic understanding of gene regulatory networks necessitates both cis- and trans-regulatory information, yet the vast majority of studies have largely focused on TF-DNA interactions, while omitting the regulatory nature of these interactions. As a result, this has precluded our ability to reconstruct the true regulatory architecture and transcriptional logic underlying gene networks. To address this fundamental gap in our knowledge, we directly and empirically measure the regulatory role of hundreds of Arabidopsis TFs, enabling the first integrated genome-scale cis- and trans-regulatory analysis to infer gene network behavior. By layering on trans-regulatory activities of TFs, we describe an entirely new approach to enrich our systems-level understanding of transcriptional regulation in any biological systems by providing directionality and causality in transcriptional networks.

Currently, most gene networks indirectly infer transcription factor regulatory activity through limited observations of TF-DNA binding and gene expression patterns, due to the lack of empirical studies that directly assay the regulatory activity of TFs. We address this shortcoming by experimentally measuring the regulatory activity of TFs on a genome-scale. Although our method is highly consistent with previous findings, there are still potential secondary mechanisms – e.g. TF phosphorylation or heterodimerization – that can alter the regulatory activity of TFs^39,40^ that our heterologous system cannot fully capture. Nonetheless, we find that the majority of our findings are validated by previous studies and *in vivo* observations. Future work focused on systematically scaling this approach to study all TFs across entire genomes will provide invaluable information to enrich our understanding of GRNs and transcriptional regulation.

Our findings add a new perspective on the complex evolution of TFs and the emergence of transcriptional wiring in biological systems. The functionality of plant TEDs in yeast suggests that core mechanisms for transcriptional activators are deeply conserved across eukaryotes. Interestingly, this trend is maintained even for TF families that have uniquely evolved in plants (i.e., the AP2-EREBP family), revealing how distinct TFs have converged upon shared biochemical features within the boundaries of universal transcriptional mechanisms. More broadly, this suggests a model where DBD of TF families may evolve somewhat independently from their TEDs, enabling extreme changes in regulatory activity in even closely related TFs. Overall, our study provides a new perspective on how to investigate the evolution of TF function, which will help reveal how complex phenotypes may have evolved.

Many important agricultural traits are dictated by TFs, thus presenting key targets for selectively modulating and engineering solutions to challenges in agriculture, bioenergy, and sustainability. Our findings provide the foundational knowledge needed to systematically map the regulatory role of TFs at a genome-scale in order to elucidate the underlying genetic wiring of plant transcriptional networks. This systems-level understanding of plants will be an invaluable resource to the broader plant biology community in understanding the circuitry and dynamics that control nearly all facets of plant physiology, development, and responses to the environment.

## Materials and Methods

### Selection of candidate TED sequences from Arabidopsis TFs

The candidate *Arabidopsis* TF sequences were obtained from the work by O’Malley *et al*^2^. The DBDs of each candidate were identified using ScanProsite^41^. The aim was to extract the longest non DBD part of the TF that could contain a TED. In case of C-or N-terminal localization, the DBD was removed from the TF sequence leaving a putative TED candidate. In case of DBD localization in the center of the protein the longest remaining TED candidate after DBD truncation was chosen. An overview of the TED and DBD localization in each TF can be found in SI Fig. 1 and the exact location of the DBD and the motif is summarized in SI Table 1.

### Construct design and assembly

The library of 529 TFs was obtained from O’Malley *et al*. and cloned into the binary vector pms7997 using Golden Gate cloning and construct specific primers (SI Table 8)^2^. Binary vector pms7997 contained the plant codon optimized GAL4 DBD (amino acid 1-147) fused to the SV40 nuclear localization signal (PKKKRKV) and the GGSGG linker peptide, connecting the DBD to individual TEDs. The synthetic TFs were driven by the MAS promoter with tNOS as a terminator. Plasmid assemblies were transformed into *E. coli* strain DH5α and purified plasmids verified via sanger sequencing using primers pms7997_insertseq_fwd & pms7997_insertseq_rev. Using this approach, we were able to clone 403 putative TEDs. The PAP1-activator fusion constructs were assembled using golden gate cloning into vector pms057 with full length PAP1 amplified from *A. thaliana* genomic DNA (Columbia ecotype). Fusions of TEDs with dCas9 were generated by replacing VP64 in vector pYPQ152 using Gibson assembly and otherwise assembled as described^30^. All vectors used for yeast experiments were generated using Gibson assembly of backbone pAI9 which contained the native yeast GAL4-DBD (amino acid 1-147) the SV40 nuclear localization signal (PKKKRKV) and the GGSGG linker peptide connecting the DBD to individual TEDs. The precise genomic locations of the native promoters NR1 and NIR1 cloned in front of GFP were chr1:29239368-29241368(-) and chr2:6808551-6810551(+) as obtained from TAIR, respectively. Full length TFs were cloned into pms057 under the control of the 35S promoter. All primers used in this study are summarized in SI Table 8. All strains and plasmid maps are available in the Inventory of Composable Elements (ICE) at https://public-registry.jbei.org.

### Agrobacterium-mediated transient transformation in *N. benthamiana*

Generated binary vectors were transformed into *A. tumefaciens* strain GV3101. Selected transformants were inoculated in liquid media with appropriate selection, for experiments diluted to OD600 = 0.5 and mixed with the assay reporter construct to a final OD_600_ = 1.0. *N. benthamiana* plants were grown for four weeks in Percival-Scientific growth chambers at 25°C in 16/8-hour light/dark cycles and 60% humidity and infiltrated as described by Sparkes *et al*.^42^. Post infiltration *N. benthamiana* plants were maintained in the same growth conditions. Leaves were harvested three days post infiltration and 16 leaf disks from two leaves per construct were collected. The leaf disks were floated on 200 µL of water in 96 well microtiter plates and GFP and RFP fluorescence measured using a BioTek Synergy H4 microplate reader (Agilent). The reporter construct for the screen was pms6370 containing GFP and dsRed expression cassettes. GFP expression was driven by a fusion of five concatenated Gal4 DNA binding sites upstream of the core WUSCHEL promoter^14^. GFP expression was normalized using dsRed driven by the MAS promoter.

### Statistical thresholds for evaluating activator and repressors populations

We established the general repressor and activator populations by first grouping all TEDs into two groups based on decreased or increased gene expression compared to the reporter only control. We then individually defined the repressor or activator population using a Kruskal-Wallis test paired with a Dunn’s ad hoc test. The first 10 consecutive TEDs leading to significantly different gene expression from the reporter measurements marked the boundaries of the populations which set the thresholds of the repressor and activator populations around −0.68 and 1.00 log_2_ fold change, respectively (SI Table 2). We further divided the activator population by defining strong activators as TEDs with log_2_ fold change >2.

### Quantification of anthocyanin content

Anthocyanin production experiments in *N. benthamiana* plants were performed as described with the divergence that the entire infiltrated leaf tissue was collected from 2 infiltrated leaves per replicate. Collected tissue was flash frozen in liquid nitrogen and freeze dried at −50 °C in vacuum for 24 h. The dried tissue was ground using bead beating for 5 min at 30 hz and 50 mg tissue was used for extraction. Anthocyanin was extracted three times using 1% hydrochloric acid in methanol and chlorophyll was removed with aqueous chloroform. Anthocyanin content was quantified by measuring absorbance at 535 nm on a Spectronic™ 200 spectrophotometer (Thermo Fisher Scientific).

### Quantitative reverse transcription PCR (RT-qPCR)

Primers targeting the GUS and Kan genes were designed using the PrimerQuest software (IDT) (Supplementary Table 7) and pre-screened for target specificity via Primer-BLAST against the *N. benthamiana* and *A. thaliana* genomes. RT-qPCR experiments were conducted on a BioRad CFX 96-well instrument using SYBR Green (BioRad). Reaction conditions were 1x ssoAdvance SYBR Green Supermix (BioRad) and 500 nM primers in 20 µL reactions, qPCR cycling parameters were 95 °C for 3 min, followed by 40 cycles of 30 s at 95 °C and 45 s at 56 °C. The linear dynamic range and efficiency of every primer set was verified over 1 × 10^2^ to 10^9^ copies per µl plasmid template. Target specificity was experimentally validated via melting temperature analysis. For total RNA isolation, ∼75 mg of leaf tissue was harvested from three plants 5 days post-infiltration, where one half of the leaf was treated with reporter alone as reference and the other half with reporter and dCas9-TED candidate as the sample. Leaf tissue was flash frozen in liquid nitrogen and RNA extracted using the EZNA Plant RNA Kit I (Omega Biotek). DNA contamination was removed by treating total RNA with Turbo DNase with inactivation reagent (Invitrogen). cDNA was generated from 1.0 µg total RNA using SuperScript IV Vilo reverse transcriptase (Thermo Fisher Scientific). RT-qPCR was carried out using 1 µl of the reverse transcription reaction as a template. For all experiments, a no template- and a no reverse transcription control was run. All primers were tested with wild type cDNA from plant tissue treated with *Agrobacterium* containing an empty vector control with Cq > 36 as the threshold for no off-target activity. The ΔΔCq method was used to determine normalized expression with GUS as the sample- and KanR as the reference gene quantified.

### Flow cytometry

For experiments in *S. cerevisiae* lab strain W303a (*MAT*a*/MATα {leu2-3,112 trp1-1 can1-100 ura3-1 ade2-1 his3-11,15} [phi+])* was used^43^. The GAL1-GFP reporter cassette was integrated into the URA3 locus. The native Gal4-TED fusions were expressed using the TEF1 promoter in a 2µ-plasmid in the reporter strain. For flow cytometry experiments all strains were grown in CSM-URA (Sunrise Science Products) media prepared following the supplier’s manual with 2% w/v Glucose, except for the positive control which was grown in 2% w/v Galactose. Experiments were performed on the BD Accuri™ C6 flow cytometer (BD Biosciences), samples were washed with cold 1x PBS (137 mmol NaCl, 2.7 mM KCl, 1.8 mM KH_2_PO_4_, 10 mM Na_2_HPO_4_) once before measurement in 1x PBS. Per sample 100,000 events were recorded and analyzed using the FlowJo™ software.

### Gene ontology enrichment of activators and TF autoregulation

DNA binding targets of TFs in this study were obtained from the Arabidopsis DAP-seq database (http://neomorph.salk.edu/PlantCistromeDB)^2^. GO term enrichment of the target genes of TFs screened in this study was performed using the g:Profiler web service accessed via the Python API with the data source limited to GO:biological process and the significance threshold method set to default (g_SCS)^44^. To study strong activators, we focused on TFs whose TEDs increased gene expression more than VP16. As many TFs target the promoters of other TFs, we excluded GO terms linked to transcription. We further reduced the amount of GO terms by only including the 10 highest enriched GO terms for the selected activator group. GO terms inside the same functional class were manually sorted.

To study autoregulation we assigned a Boolean value to every TF, whose TED we studied here, based on whether it had been previously reported to bind its own promoter region. The Boolean value 1 was assigned to TFs with binding and 0 to TFs with no binding to their own promoter region. We then grouped the Boolean values into the TED populations and studied a potential auto-regulation bias between TED populations using a two-sided Fisher’s exact test.

### Generating a trans-regulatory annotated nitrogen response GRN

The extended nitrogen response GRN was built on a version including DNA binding information and a co-expression machine learning model based on temporal RNA-seq data^8^. In short, Varala *et al*. performed a temporal RNA-seq study of the response of *A. thaliana* to nitrogen feeding after nitrogen starvation. They collected tissue of plants at different time points post nitrogen feeding as well as an untreated control. By comparing the RNA-seq profile of the induced and untreated samples they established a set of nitrogen responsive genes, defined as the first time point where expression in the induced sample was ≥ 1.5 fold of control. They further generated a pruned GRN only including TF-Gene interaction that had previously been verified *in vitro*. We utilized this network to annotate TF-Gene interactions with TED data, represented as an edge attribute. We visualized the GRN and extracted TF-gene interactions of NR1 and NIR1 using Cytoscape v3.9.0^45^. To study the effects of the activator group observed at 15 min we performed RNA-seq analysis using the limma package and DESeq2 in R as shown in Varala *et al*.^46,47^. Rate of induction of gene expression was calculated by subtracting fold changes of consecutive time points. The time point of maximal induction and maximal decrease of gene expression was derived from these subtractions.

### Validation of GRN predicted TF-gene interactions

To verify regulatory activity of TFs binding NR1 and NIR1 promoter regions in the nitrogen GRN, we utilized *Agrobacterium* mediated transient transformation as described above. For each condition we used three *Agrobacterium* strains: one expressing a native full-length TF, one with a GFP reporter driven by the respective promoter regions of NR1 or NIR1 and one with mScarlet driven by the NOS promoter for normalization. Modulation of gene expression from basal levels was validated using the Mann-Whitney-U test.

### Localization of activation domains in plant TFs using ADpred

We localized putative activation domains in the TEDs from our study using the ADpred model^36^. The model analyzes sequence stretches of 30 amino acids with predicted secondary structure information. Therefore, the secondary structure of full-length TED domains was predicted using the PsiPred workbench^48^. For the analysis of individual TEDs, we fragmented the protein sequence and secondary structure prediction into 30 amino acid sections moving 5 amino acids in between fragments. If any of the fragments of a given TED scored at ≥ 0.9 in the ADpred model the TED potentially contained an AD. We assigned a Boolean to every TED based on the scoring, 0 for no AD and 1 for containing a potential AD. A potential bias between different TED populations was observed using a two-sided Fisher’s exact test. To validate the predictive power of the model we chose 3 strong activators and subcloned the longest consecutive amino acid stretches scoring above the ADpred threshold of 0.9 into pms7997 and assayed their performance in comparison to their TED counterparts in *N. benthamiana*.

## Acknowledgements

We thank Ronan O’Malley for providing template DNA for the amplification of all TED regions of interest and both Simon Alamos and James Nunez for reviewing the manuscript.

## Funding

This work was part of the DOE Joint BioEnergy Institute (http://www.jbei.org) supported by the U. S. Department of Energy, Office of Science, Office of Biological and Environmental Research through contract DE-AC02-05CH11231 between Lawrence Berkeley National Laboratory and the U.S. Department of Energy. The United States Government retains and the publisher, by accepting the article for publication, acknowledges that the United States Government retains a non-exclusive, paid-up, irrevocable, worldwide license to publish or reproduce the published form of this manuscript, or allow others to do so, for United States Government purposes.

## Author contributions

Project design: NFCH, BL, PMS

Experimental work: NFCH, AZ, BL, KM, IJO

Analyses: NFCH, AZ

Visualization: NFCH, AZ

Financial support: PMS

Supervision: PMS

Writing – original draft: NFCH, AZ, PMS

Writing – review & editing: NFCH, AZ, KM, PMS

## Competing interests

PMS and NFCH have a patent pending for the library of synthetic TFs and all derived parts. The remaining authors have no conflict of interest to declare.

## Data and materials availability

All plasmid materials and bacterial strains will be made available through the Inventory of Composable Elements (https://registry.jbei.org/). Sequences and raw data are available as supplementary materials.

## Notes

### Competing Interest Statement

The authors have declared no competing interest.

