## Supplementary Information for "The trans-regulatory landscape of gene networks in plants"

**This PDF file includes:**

Figs. S1 to S8

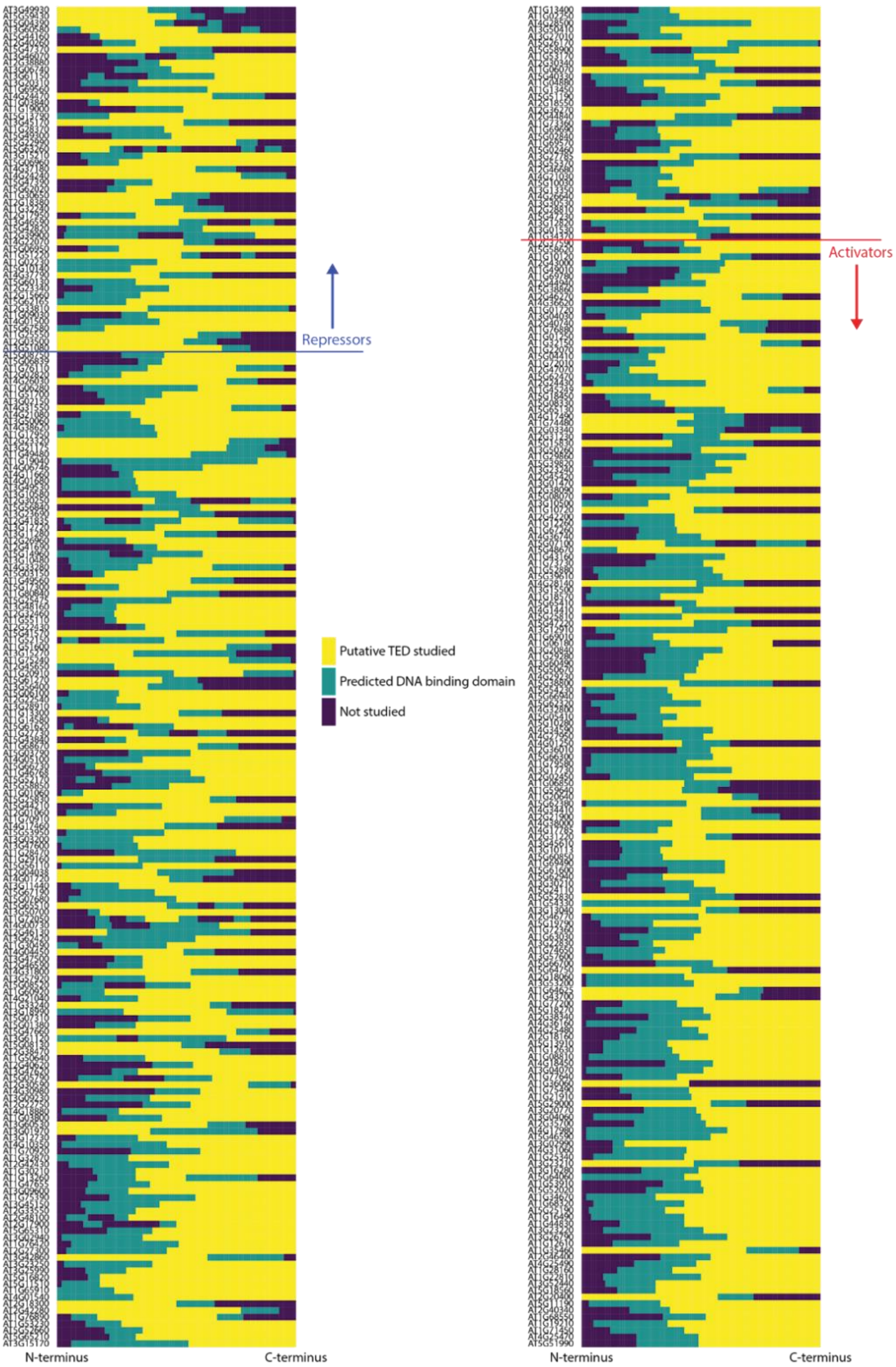


**Figure S1. Overview of extracted putative TED regions for all TFs studied normalized to the full length of each protein.** TEDs above the blue line belong to the repressor population, below the red line to activators. Yellow: Putative TED studied, Green: Predicted DBD, Blue: Regions not studied in this work.

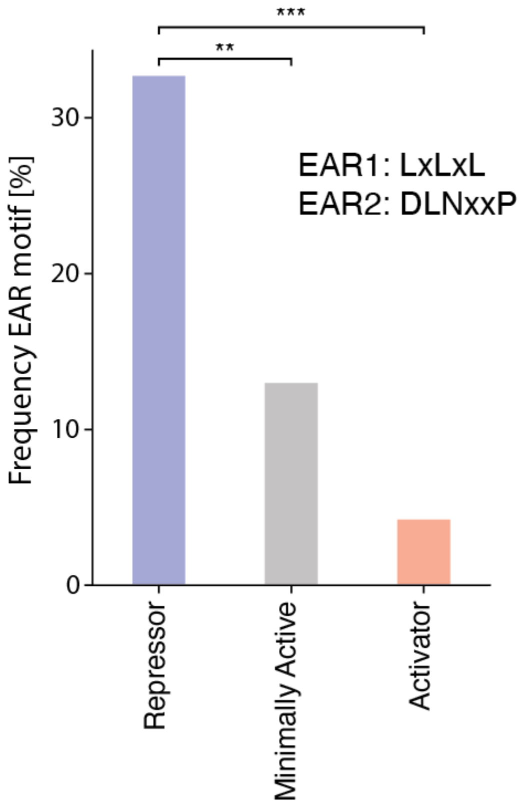


**Figure S2. The repressive EAR motifs are enriched in the repressor population.** A Boolean value was associated to every TED based on the occurrence of EAR motifs (LxLxL or DLNxxP) in their protein sequence. Statistical differences between populations were verified by a two-sided Fisher's exact test. Asterisks indicate results of Fisher's exact test ** $P \leq 5 \times 10^{-3}$, *** $P \leq 5 \times 10^{-4}$.

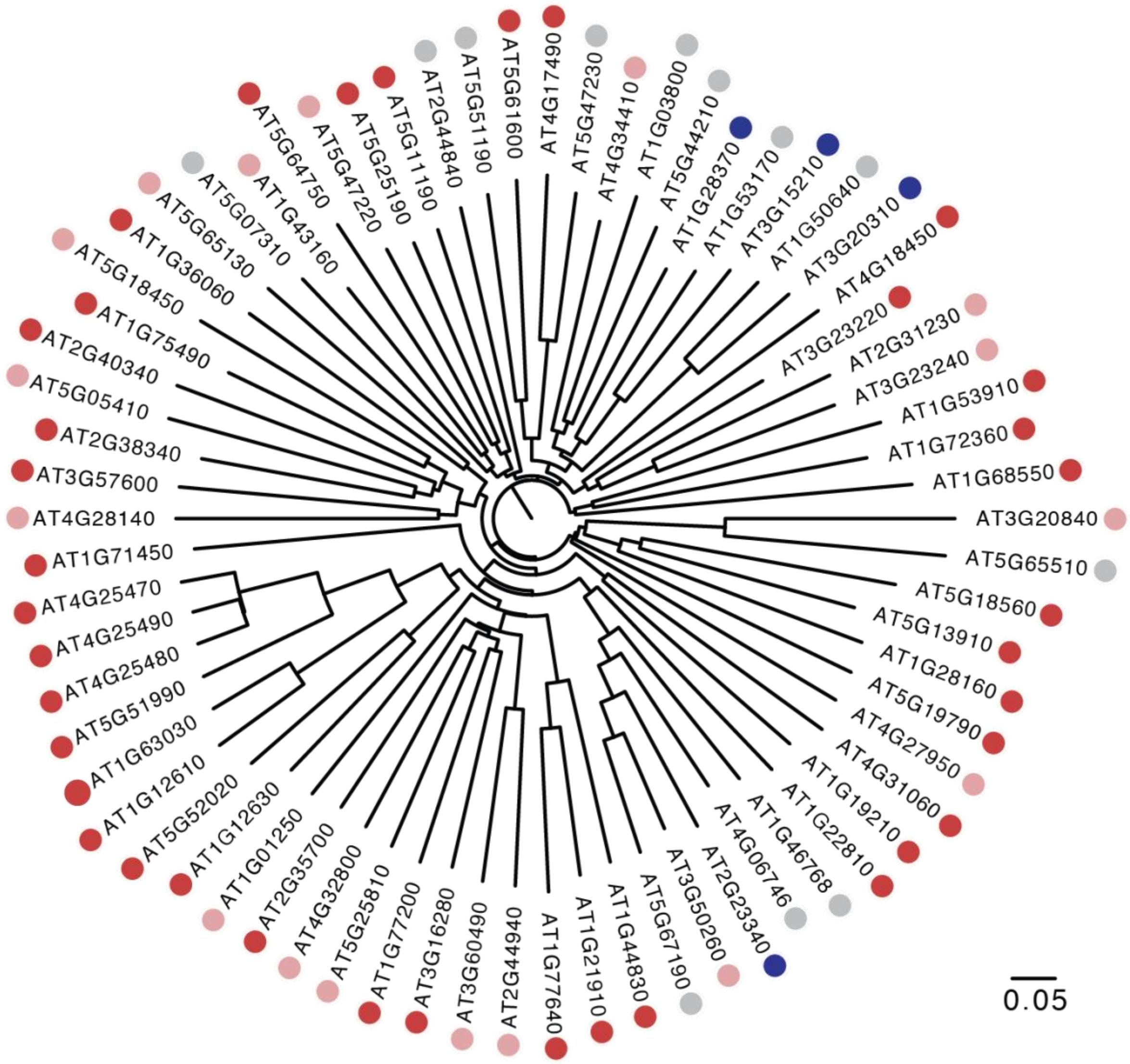


**Figure S3. Homology based phylogeny of all TFs of the AP2-EREBP TF family with TEDs characterized in this work.** Colored dots indicate activity with blue: repressors, gray: minimally active, pink: activators, red: strong activators ($log_2$ Fold change > 2). Scale bar indicates phylogenetic distance based on sequence homology from multiple sequence alignment.

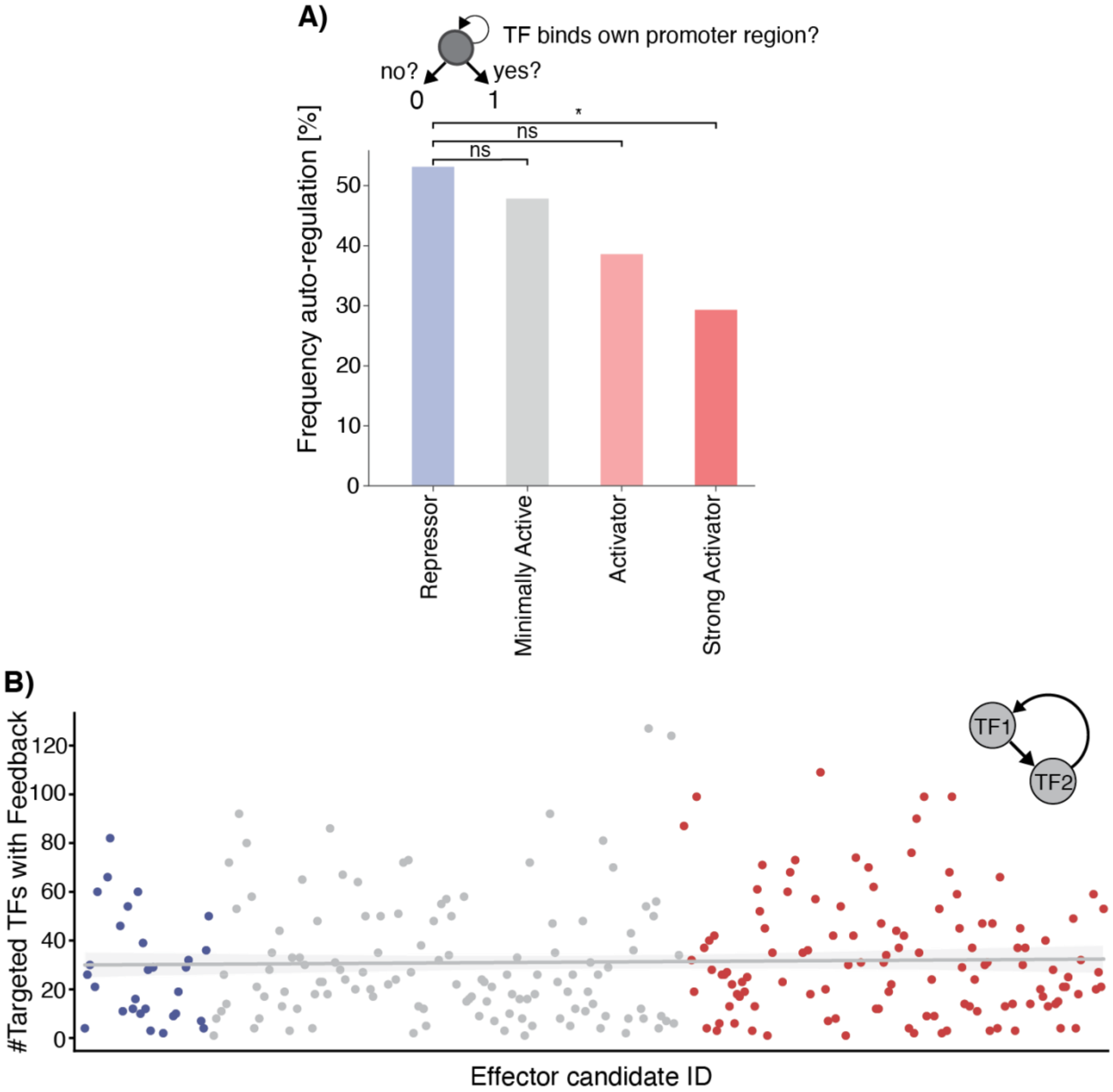


**Figure S4. Repressors are more likely to autoregulate their own expression than strong activators.** (**A**) A Boolean value was associated to every TF with characterized TED based on their potential to bind their own promoter region. Biases between TED populations were verified by a two sided Fisher's exact test ( * $P \leq 5 \times 10^{-2}$). (**B**) There is no observable trend for feedback loops between TED populations. Sum of targeted TFs binding the initial TFs promoter region. Blue dots represent repressors, gray dots minimally active TEDs and red dots activators.

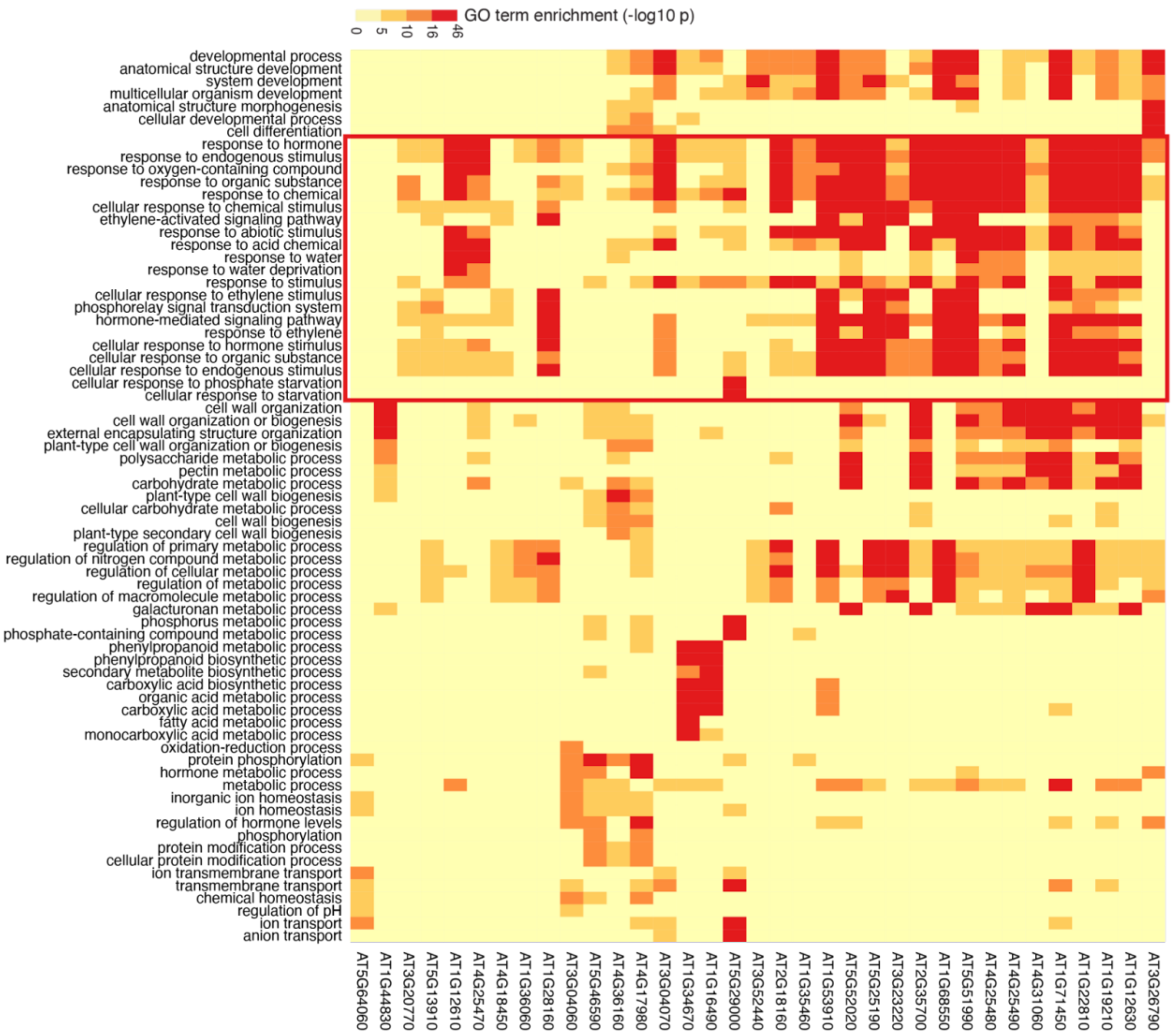


**Figure S5. Genomic targets of strong activators are enriched for GO terms linked to the response to stimuli.** Gene ontology enrichment for biological process terms of TF targeted genes were generated using GProfiler and GO terms linked to transcription omitted to exclude overrepresentation of TFs targeted by the activator group. The 10 most enriched GO terms for each activator are included. Red box indicates GO terms linked to the response to stimuli.

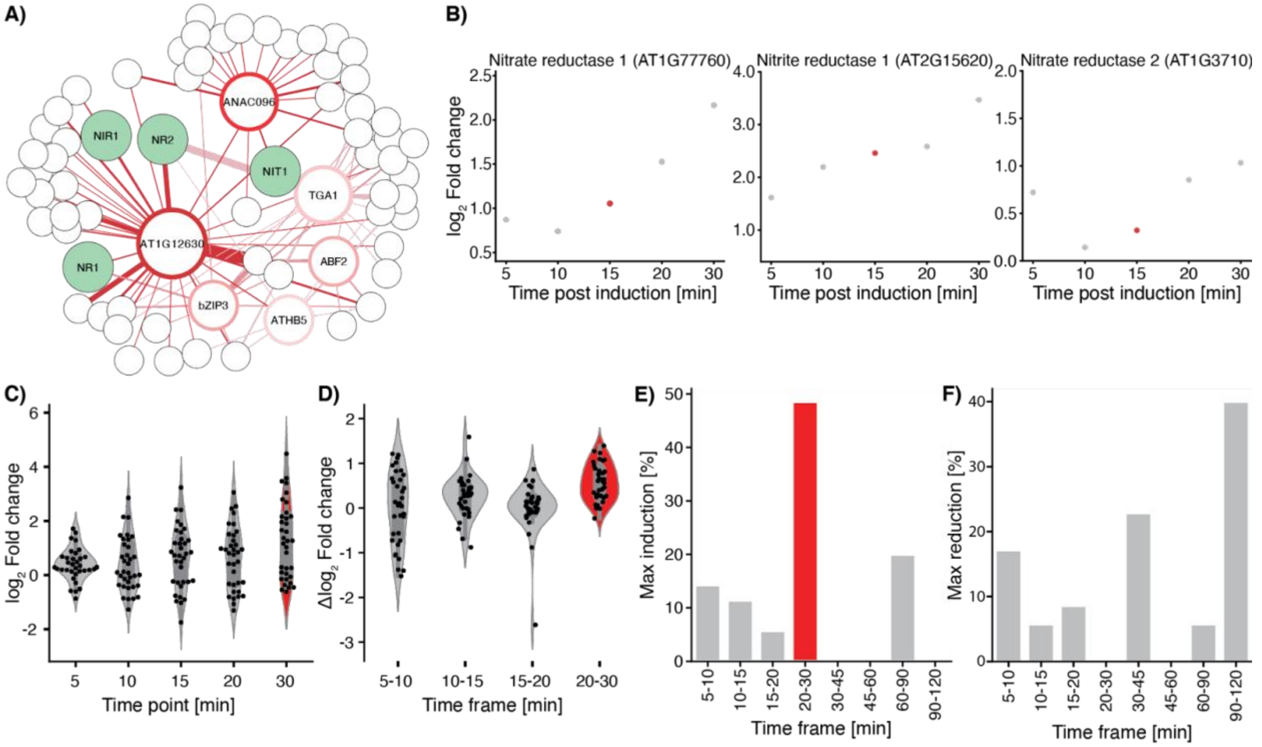


**Figure S6. Expression of TF activators at 15 min during Arabidopsis nitrate response results in the activation of downstream genes.** (**A**) Subnetwork derived from (A) with induced TFs at 15 min post nitrate induction and their respective genomic targets. (**B**) Time course of gene expression of nitrate reductase 1, nitrite reductase 1 and nitrate reductase 2. The time point where the cluster of activators from Fig. 2B shows significant increases in RNA abundance is marked in red. (**C**) Expression profiles of genes targeted by induced TFs at 15 min in comparison to their respective expression at 0 min. Fold changes were calculated using a multiple variable linear model comparing both non induced sample and time point 0. Each dot represents one gene. (**D**) Distributions for the rate of expression change between timepoints for the genes in (B). Rates are calculated by subtracting the fold change expression of the previous time point from the current for every gene. (**E**) Based on (C) fraction of genes showing time step with largest rate of gene expression induction throughout the entire time course. (**F**) Based on (C) fraction of genes showing time step with largest rate of gene expression reduction throughout the entire time course.

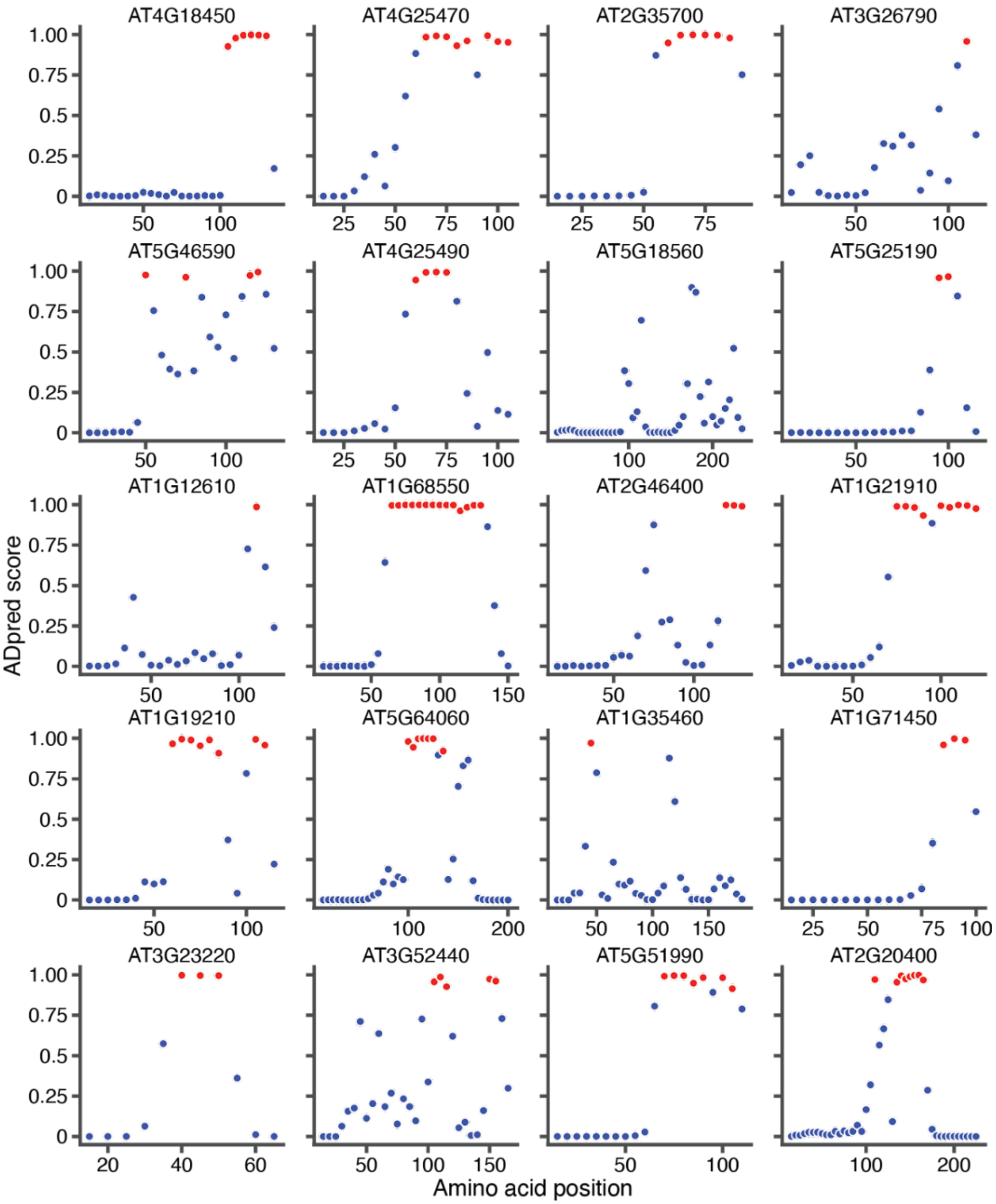


**Figure S7. ADpred predicts putative activation domains in plant TFs.** ADpred evaluation of 20 activators stronger than VP16. ADpred scores were calculated for every 30 amino acid stretch slid along the protein sequence with window size = 5. Red dots indicate 30 amino acid long segments with ADpred score ≥ 0.9, blue dots < 0.9.

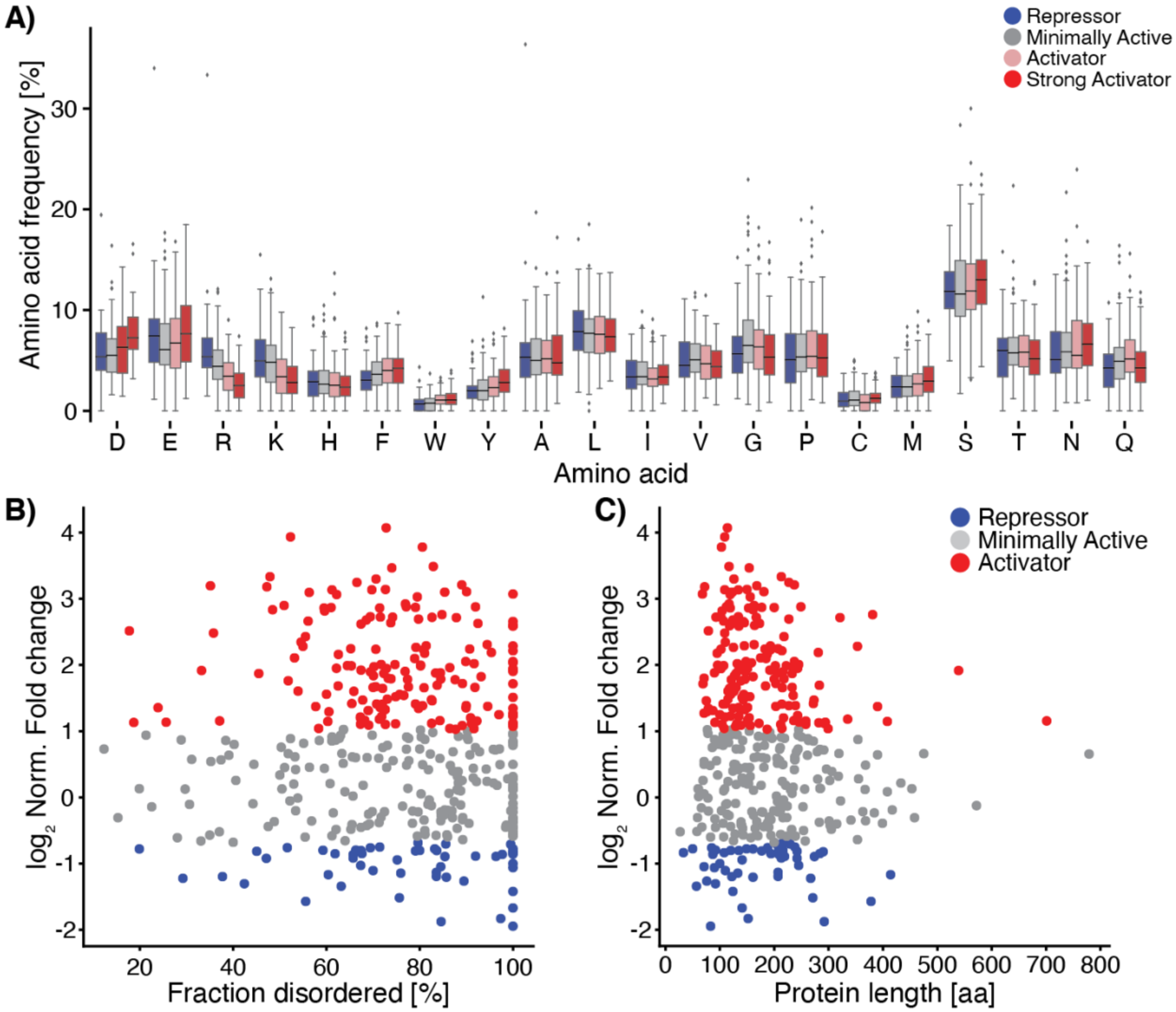


**Figure S8. TED populations only show differences in distinct biochemical features.** (**A**) Distribution of individual amino acid frequency inside each TED population. (**B**) Fraction of protein sequence predicted to be disordered by VSL2 in relation to normalized GFP fold change. (**C**) Length of every TEDs protein sequence in relation to normalized GFP fold change.
